# The EARP Complex and its Interactor EIPR-1 are Required for Cargo Sorting to Dense-Core Vesicles

**DOI:** 10.1101/046003

**Authors:** Irini Topalidou, Jérôme Cattin-Ortolá, Andrea L. Pappas, Kirsten Cooper, Gennifer E. Merrihew, Michael J. MacCoss, Michael Ailion

## Abstract

The dense-core vesicle is a secretory organelle that mediates the regulated release of peptide hormones, growth factors, and biogenic amines. Dense-core vesicles originate from the trans-Golgi of neurons and neuroendocrine cells, but it is unclear how this specialized organelle is formed and acquires its specific cargos. To identify proteins that act in dense-core vesicle biogenesis, we performed a forward genetic screen in *Caenorhabditis elegans* for mutants defective in dense-core vesicle function. We previously reported the identification of two conserved proteins that interact with the small GTPase RAB-2 to control normal dense-core vesicle cargo-sorting. Here we identify several additional conserved factors important for dense-core vesicle cargo sorting: the WD40 domain protein EIPR-1 and the endosome-associated recycling protein (EARP) complex. By assaying behavior and the trafficking of dense-core vesicle cargos, we show that mutants that lack EIPR-1 or EARP have defects in dense-core vesicle cargo-sorting similar to those of mutants in the RAB-2 pathway. Genetic epistasis data indicate that RAB-2, EIPR-1 and EARP function in a common pathway. In addition, using a proteomic approach in rat insulinoma cells, we show that EIPR-1 physically interacts with the EARP complex. Our data suggest that EIPR-1 is a new component of the EARP complex and that dense-core vesicle cargo sorting depends on the EARP-dependent retrieval of cargo from an endosomal sorting compartment.

**Author Summary:** Animal cells package and store many important signaling molecules in specialized compartments called dense-core vesicles. Molecules stored in dense-core vesicles include peptide hormones like insulin and small molecule neurotransmitters like dopamine. Defects in the release of these compounds can lead to a wide range of metabolic and mental disorders in humans, including diabetes, depression, and drug addiction. However, it is not well understood how dense-core vesicles are formed in cells and package the appropriate molecules. Here we use a genetic screen in the microscopic worm *C. elegans* to identify proteins that are important for early steps in the generation of dense-core vesicles, such as packaging the correct molecular cargos in the vesicles. We identify several factors that are conserved between worms and humans and point to a new role for a protein complex that had previously been shown to be important for controlling trafficking in other cellular compartments. The identification of this complex suggests new cellular trafficking events that may be important for the generation of dense-core vesicles.

## Introduction

The dense-core vesicle is a specialized secretory organelle found in neurons and endocrine cells. In endocrine cells, these vesicles are often called secretory granules; for simplicity, we will here use the term ‘dense-core vesicles’ to refer to both the neuronal and endocrine carriers. Dense-core vesicles package several classes of cargo, including peptide hormones, growth factors, and biogenic amines. In response to elevated calcium, dense-core vesicles release their cargos to modulate a variety of biological processes ranging from blood glucose homeostasis to development, function and plasticity of the nervous system. Despite the importance of the signaling molecules released by dense-core vesicles, surprisingly little is known about how dense-core vesicles are generated and acquire their proper cargos (1–4).

Dense-core vesicles originate at the trans-Golgi network, but it is unclear how their compartmental identity is established and how cargos are correctly sorted into nascent dense-core vesicles. After budding from the trans-Golgi, immature dense-core vesicles undergo a poorly defined maturation process that involves homotypic vesicle fusion, peptide processing, and the removal of some soluble and transmembrane cargos (5–11). Some studies suggest that luminal dense-core vesicle cargos sort purely through an intrinsic ability to aggregate in the low pH/high Ca^2+^ milieu of the trans-Golgi (12), while other studies suggest that aggregation is not sufficient and that sorting relies on poorly defined structural motifs in cargo proteins that may interact with multiple sorting receptors (13,14). Two general models of dense-core vesicle cargo sorting have been debated for more than twenty years: the ‘sorting by entry’ and ‘sorting by retention’ models that propose that sorting occurs in the trans-Golgi as dense-core vesicles bud off, or in a post-Golgi step where non-dense-core vesicle cargos are removed (15,16). Experimental evidence supports both models, which are not mutually exclusive, suggesting that both mechanisms contribute to dense-core vesicle biogenesis.

Molecular mechanisms of dense-core vesicle biogenesis are poorly understood largely because few proteins have been identified that function in this process. In recent years, the small GTPase Rab2 has emerged as a major regulator of dense-core vesicle cargo trafficking. RAB-2, its effector ICA69/RIC-19, and the putative RAB-2 GTPase-activating protein (GAP) TBC-8 have all been shown to be important for early steps in dense-core vesicle cargo sorting at or near the trans-Golgi (17–22). To identify additional factors in this pathway, we used a forward genetic screen in the nematode *C. elegans* to isolate mutants that affect dense-core vesicle function. We identified RUND-1, a RUN domain protein, and CCCP-1, a coiled-coil protein, that act as RAB-2 effectors to mediate normal dense-core vesicle cargo sorting (23). In mutants of *rab-2*, *rund-1*, or *cccp-1*, morphologically normal dense-core vesicles are generated and transported to their axonal release sites, but these dense-core vesicles have reduced levels of both luminal and transmembrane cargos. RAB-2 and its effectors colocalize at or near the trans-Golgi, suggesting that they act in the neuron cell body to mediate proper cargo-sorting as dense-core vesicles bud off from the trans-Golgi or undergo post-Golgi maturation.

Here we investigate the function of several additional molecules identified in our genetic screen for dense-core vesicle mutants: the conserved WD40 domain protein EIPR-1 and the VPS-52 and VPS-53 subunits shared by the Golgi-associated retrograde protein (GARP) and endosome-associated recycling protein (EARP) complexes. We show that EIPR-1 physically and genetically interacts with the EARP complex. Mutants that lack EIPR-1 or EARP have defects in dense-core vesicle cargo-sorting similar to those of mutants that lack RAB-2, RUND-1 or CCCP-1, and genetic epistasis data indicate that they function in the same pathway. The requirement of EARP for proper sorting of cargos to dense-core vesicles suggests that dense-core vesicle cargo sorting is achieved in part through the retrieval of dense-core vesicle cargos or cargo receptors from an endosomal sorting compartment. While there is ample prior evidence that cargos are removed during dense-core vesicle maturation and trafficked to endosomes, our experiments suggest that there is also trafficking back from endosomes to maturing dense-core vesicles.

## Results

### *eipr-1* encodes a conserved WD40 domain protein

To identify proteins required for dense-core vesicle biogenesis, we performed a forward genetic screen in the nematode *C. elegans* based on suppression of the hyperactive locomotion of an activated Gq mutant (23). We previously characterized the function of two novel conserved proteins identified from this screen, RUND-1 and CCCP-1. Both RUND-1 and CCCP-1 interact with the small GTP-binding protein RAB-2 when it is in its activated GTP-bound state (23). Mutants in *rab-2*, *rund-1* and *cccp-1* have a stereotypical defect in locomotion behavior characterized by little spontaneous movement on food, but coordinated slow locomotion when stimulated. One additional uncloned mutant from our screen (*ox316*) had defects in locomotion and egg-laying very similar to *rab-2*, *rund-1*, and *cccp-1* mutants. We cloned this mutant by single nucleotide polymorphism (SNP) mapping and transgenic rescue experiments (Materials and Methods). We named the gene *eipr-1* (<u>E</u>ARP-<u>i</u>nteracting-<u>pr</u>otein, see below). The *eipr-1* mutant phenotype is fully rescued by microinjection of the single gene Y87G2A.11, and the *ox316* mutation leads to a premature stop codon in this gene (Figure 1A-C). We subsequently obtained a deletion allele (*tm4790*) that causes identical locomotion and egg-laying phenotypes (Figure 1C; Figure S1A). *eipr-1* encodes a novel WD40 domain protein of 362 amino acids (Figure 1B). Like RUND-1 and CCCP-1, EIPR-1 has a single conserved ortholog in other metazoans. However, unlike RUND-1 and CCCP-1, EIPR-1 is also found outside metazoans in plants and some fungi and protozoa (Figure S2). Interestingly, EIPR-1 is not present in all protozoa and fungi, suggesting an ancestral origin and loss in specific lineages including the budding yeast *Saccahromyces cerevisiae*. The mouse *Eipr1* cDNA could also rescue the worm *eipr-1* mutant (Figure 1C; Figures S1A), demonstrating functional conservation of the mammalian ortholog.

**Figure 1.**
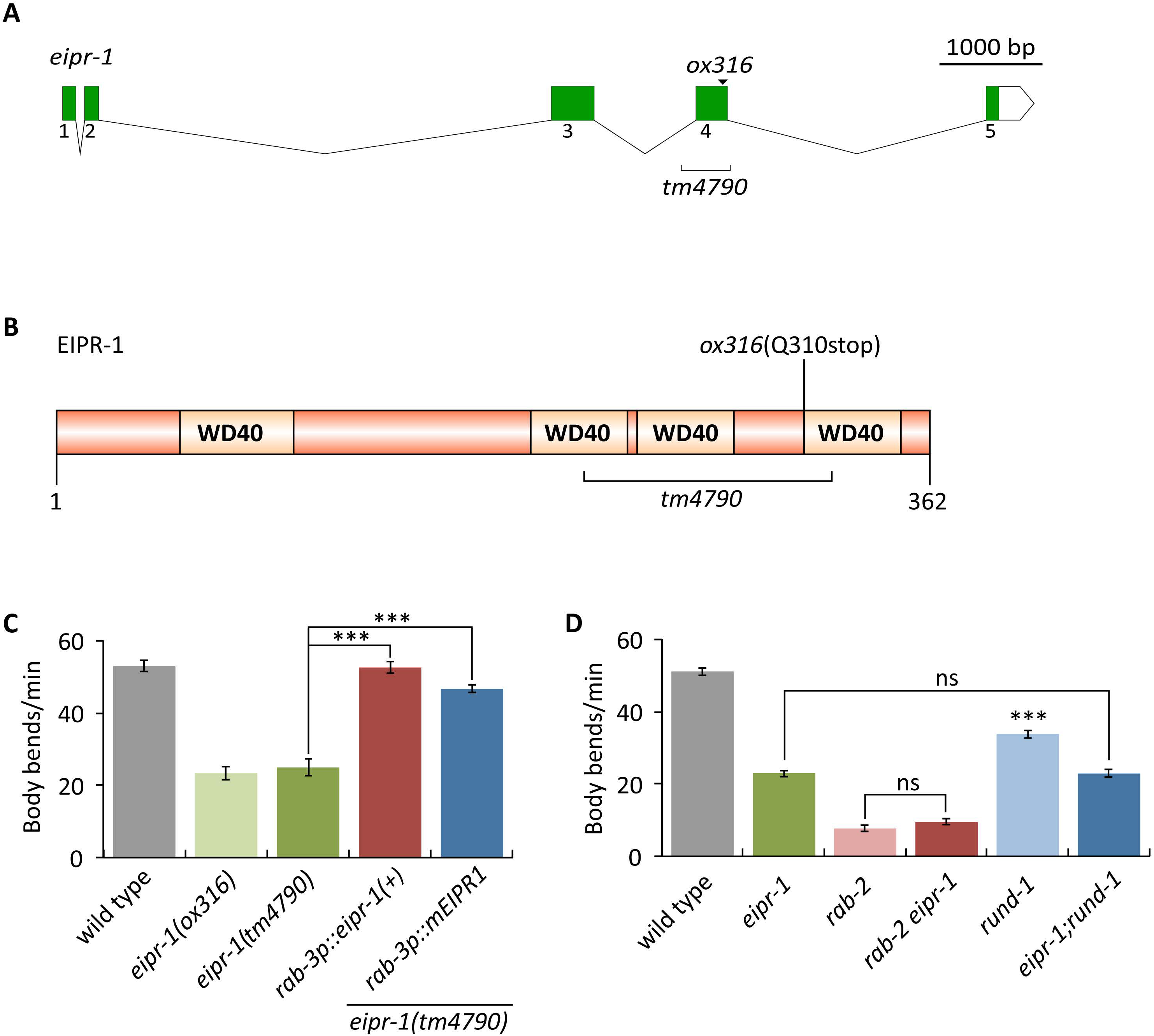
*eipr-1* encodes a WD40 domain protein that controls locomotion behavior. (A) Gene structure of *eipr-1*. Solid boxes show coding segments. A white box shows the 3’ untranslated region. The positions of the *ox316* stop mutation and the *tm4790* deletion are shown. The gene structures were drawn using Exon-Intron Graphic Maker (http://wormweb.org/exonintron). (B) Domain structure of the EIPR-1 protein. EIPR-1 is a 362 amino acid protein with four predicted WD40 repeats. The part of the protein deleted by *tm4790* is marked, but because this deletion starts and ends in introns (Figure 1A), its precise effect on the protein is unknown, though it would cause a frameshift if exon 3 splices to exon 5 in this mutant. The protein structure was drawn using DOG 1.0 (Ren et al., 2009). WD40 repeat positions were as predicted by SMART (http://smart.embl-heidelberg.de/). (C) *eipr-1*acts in neurons to control locomotion. The *eipr-1* mutant has slow locomotion that is rescued by panneuronal expression of either the worm gene or its mouse ortholog (***, P<0.001). Error bars = SEM; n = 10. (D) *eipr-1* acts in the same genetic pathway as *rab-2* and *rund-1* to control locomotion. Double mutants of *eipr-1* with *rab-2* or *rund-1* do not have stronger locomotion defects than the single mutants (***, P<0.001 compared to wild type; ns, not significant, P>0.05). Error bars = SEM; n = 10.

### EIPR-1 acts in the RAB-2 pathway to control locomotion and dense-core vesicle cargo trafficking

The phenotypic similarities between *eipr-1* and *rab-2*, *rund-1*, and *cccp-1* mutants suggests that these genes act in a common pathway. To test this, we built double mutants between *eipr-1* and either *rab-2* or *rund-1* and assayed locomotion and egg-laying behavior. In both cases, the double mutants do not show enhanced locomotion or egg-laying defects compared to the single mutants (Figure 1D; Figure S1B). Thus, *eipr-1* acts in the *rab-2* pathway to control locomotion and egg-laying behaviors.

Mutants in *rab-2*, *rund-1* and *cccp-1* have reduced axonal levels of dense-core vesicle cargos (19,22,23). To test whether *eipr-1* also acts in dense-core vesicle cargo trafficking, we examined trafficking of both luminal and membrane cargos of dense-core vesicles. For luminal cargos, we assayed the Venus-tagged peptides NLP-21, FLP-3, and INS-22. NLP-21 and FLP-3 are endogenous neuropeptides that undergo proteolytic processing (24,25), while INS-22 is an insulin-family peptide that does not undergo proteolytic processing (26). For a membrane cargo, we assayed GFP tagged IDA-1, the worm ortholog of the dense-core vesicle transmembrane protein phogrin (27). Like the *rab-2* pathway mutants, we found that *eipr-1* mutants also had reduced axonal levels of all luminal cargos and the transmembrane cargo IDA-1::GFP (Figure 2A-D; Figure S3A-C). Double mutants of *eipr-1* with either *rab-2* or *rund-1* did not show an enhanced defect compared to the single mutants (Figure 2A), again suggesting that *eipr-1* acts in the *rab-2* pathway. To determine whether the *eipr-1* defect is specific to dense-core vesicle cargos, we examined trafficking of the synaptic vesicle cargo UNC-17, the vesicular acetylcholine transporter.*eipr-1* mutants showed no reduction in axonal levels of UNC-17 (Figure 2E). *eipr-1* mutants also showed no reduction in axonal levels of the synaptic vesicle SNARE protein synaptobrevin (SNB-1) (Figure S3D). We conclude that EIPR-1 acts in the RAB-2 pathway to specifically mediate trafficking of dense-core vesicle cargos.

**Figure 2.**
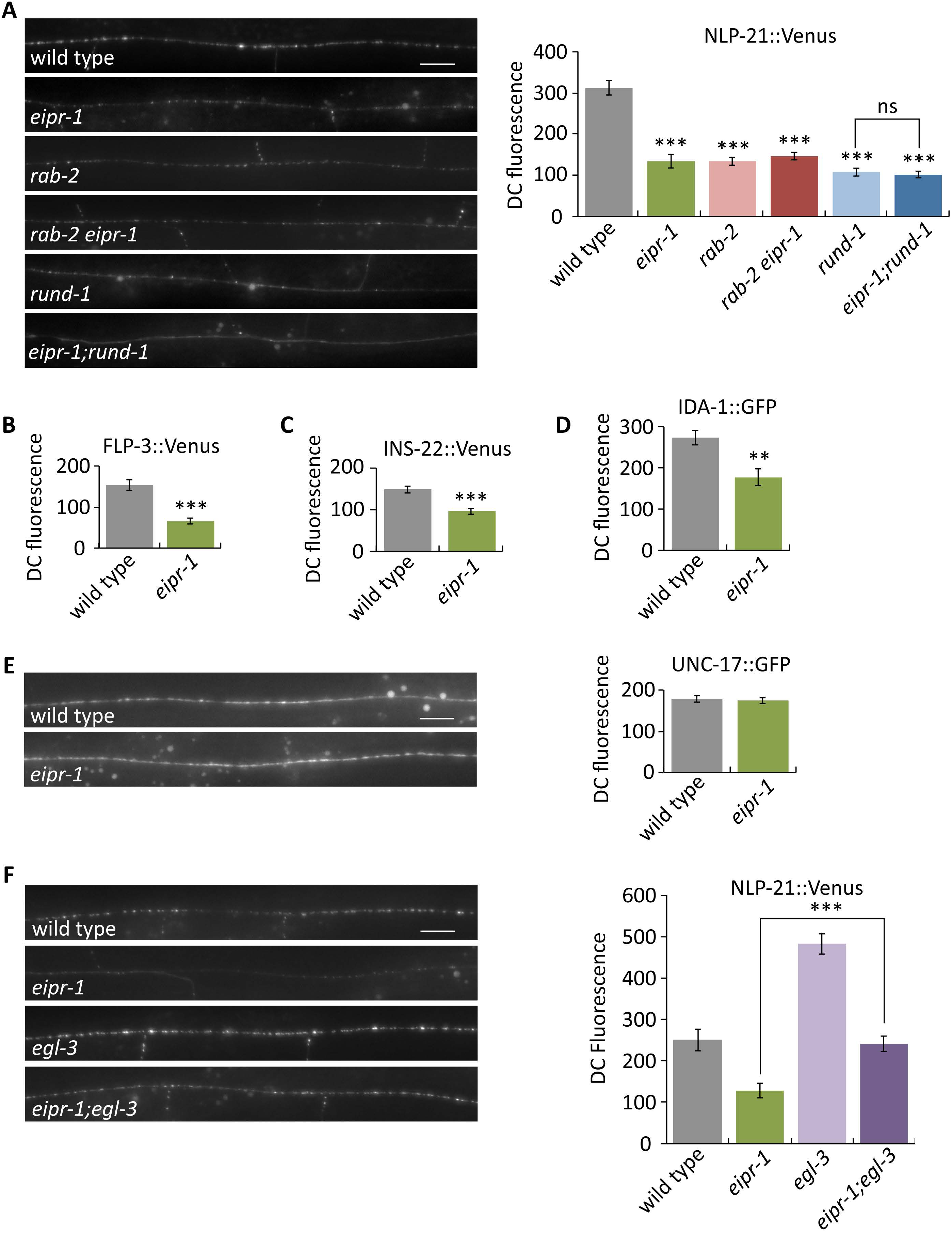
*eipr-1* mutants have defects in trafficking dense-core vesicle cargos. (A) Left: representative images of NLP-21::Venus fluorescence in motor neuron axons of the dorsal nerve cord. Scale bar: 10 μm. Right: quantification of NLP-21::Venus fluorescence levels in the dorsal nerve cord. The mean fluorescence intensity is given in arbitrary units. *eipr-1*, *rab-2*, and *rund-1* mutants have decreased levels of fluorescence in the dorsal cord (DC), indicative of an NLP-21::Venus sorting or trafficking defect (***, P<0.001 compared to wild type). Double mutants of *eipr-1* with *rab-2* or *rund-1* are not significantly different than the single mutants (P>0.05). Error bars = SEM; n = 10. (B) FLP-3::Venus fluorescence levels in the dorsal nerve cord. ***, P<0.001. Error bars = SEM; n = 12. (C) INS-22::Venus fluorescence levels in the dorsal nerve cord. ***, P<0.001. Error bars = SEM; n = 11-12. (D) IDA-1::GFP fluorescence levels in the dorsal nerve cord. **, P<0.01. Error bars = SEM; n = 13-14. (E) UNC-17::GFP fluorescence levels in the dorsal nerve cord. *eipr-1* mutants do not have a defect (P>0.05). Scale bar: 10 μm. Error bars = SEM; n = 16-19. (F) The peptide processing mutant *egl-3* increases the level of NLP-21::Venus in the dorsal cord in both wild type and *eipr-1* mutant backgrounds. Left: representative images. Scale bar: 10 μm. Right: quantification of NLP-21::Venus fluorescence levels in the dorsal nerve cord. ***, P<0.001. Error bars = SEM; n = 9-12.

A mutation in the *egl-3* proprotein convertase blocks neuropeptide processing and leads to an increase in axonal levels of luminal dense-core vesicle cargos (19,22). As with *rab-2* and *rund-1*, there is increased axonal fluorescence of Venus-tagged NLP-21 in *eipr-1*; *egl-3* double mutants as compared to the *eipr-1* single mutant (Figure 2F). However, the *egl-3* mutant on its own has increased axonal fluorescence compared to wild-type, and the *eipr-1* mutant decreases axonal fluorescence about 50% in either wild-type or *egl-3* backgrounds (Figure 2F), indicating that *eipr-1*, like *rab-2* and *rund-1*, acts in parallel to *egl-3*. Thus, *eipr-1* mutants have reduced trafficking of both processed and unprocessed cargos.

### EIPR-1 acts in neurons to control locomotion and dense-core vesicle biogenesis

The behavioral and dense-core vesicle phenotypes of *eipr-1* mutants suggest that it acts in neurons. Given that *eipr-1* is the fourth gene in an operon, it has been difficult to determine the endogenous expression pattern of *eipr-1* (see Materials and Methods for details and Figure S4A). Thus, we instead expressed the *eipr-1* cDNA under the control of neuronal-specific promoters to determine whether the gene functions in neurons. Expression of *eipr-1(+)* under the neuronal specific *rab-3* promoter fully rescued *eipr-1* mutant locomotion, dense-core vesicle trafficking, and egg-laying (Figure 3A,B and data not shown). Expression of *eipr-1(+)* under the cholinergic *unc-17* promoter also rescued the locomotion defect. Expression in cholinergic motor neurons (using the *acr-2* promoter) did not rescue the locomotion phenotype, but expression driven by a head-specific derivative of the *unc-17* promoter *(unc-17Hp)* rescued the locomotion phenotype, indicating that *eipr-1* acts mainly in head cholinergic neurons to regulate locomotion behavior (Figure 3A). The dense-core vesicle trafficking defect of *eipr-1* mutants is cell autonomous, as expression of *eipr-1(+)* driven by the *unc-129* promoter fully rescued the axonal trafficking defect of NLP-21::Venus expressed by the same *unc-129* promoter (Figure 3B). Thus, EIPR-1 acts in neurons to control dense-core vesicle trafficking at the cellular level and locomotion behavior at the organismal level.

**Figure 3.**
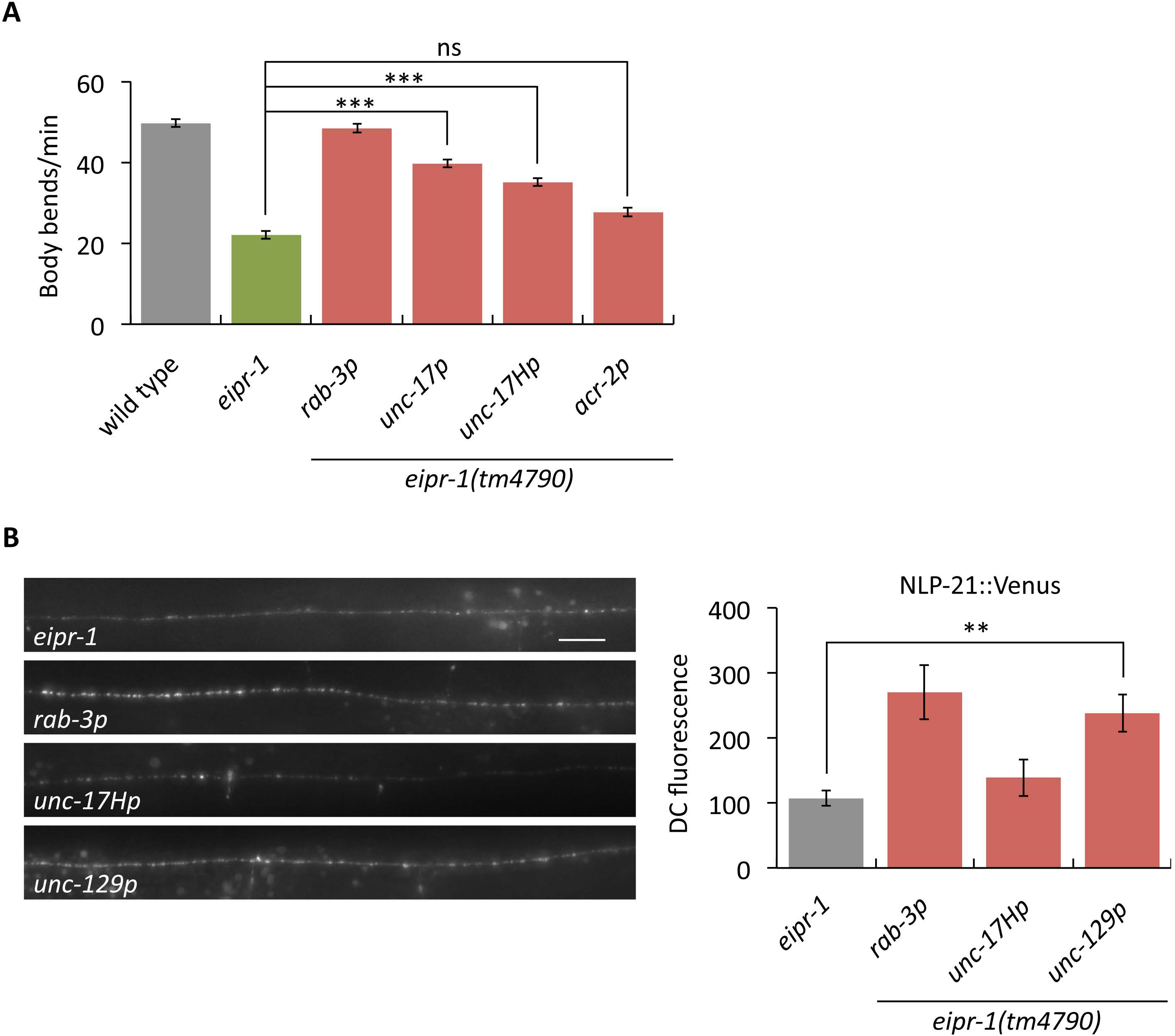
*eipr-1* acts in head cholinergic neurons to control locomotion. (A) The *eipr-1*cDNA was expressed in an *eipr-1* mutant background using the following promoters: *rab-3*(all neurons), *unc-17* (cholinergic neurons), *unc-17H* (a derivative of the *unc-17* promoter that lacks the enhancer for ventral cord expression and thus expresses only in head cholinergic neurons), and *acr-2* (ventral cord cholinergic motor neurons). Expression driven by the *rab-3, unc-17*, and *unc-17H* promoters rescued the mutant locomotion defect, but expression driven by *acr-2* did not (***, P<0.001; ns, not significant, P>0.05). Error bars = SEM; n = 12-20. (B) *eipr-1* acts cell autonomously to control dense-core vesicle cargo trafficking. The *eipr-1*cDNA was expressed in an *eipr-1* mutant background using either the *rab-3* promoter (all neurons) or *unc-129* promoter (a subset of cholinergic motor neurons where NLP-21::Venus is also expressed). Expression driven by both promoters rescued the unc-129p::NLP-21::Venus trafficking defect of the mutant (***, P<0.001). Left: representative images. Scale bar: 10 μm. Right: quantification of dorsal cord fluorescence. Error bars = SEM; n = 7-13.

The cell-specific constructs described above had EIPR-1 C-terminally tagged with GFP. Because these constructs were able to rescue the mutant phenotype, the EIPR-1::GFP protein must be functional; thus we examined these strains to determine the subcellular localization of EIPR-1. Unlike RAB-2, RUND-1 and CCCP-1 which are localized predominantly to the trans-Golgi (19,22,23), EIPR-1::GFP was expressed diffusely throughout the cytoplasm when expressed under such promoters as *rab-3* or *unc-17*, even when coexpressed with a GTP-locked mutant of RAB-2 (Figure S4B,C). However, given that endogenously tagged EIPR-1 is undetectable (see Materials and Methods), these EIPR-1::GFP constructs are almost certainly overexpressed even though they are single-copy insertions, and the lack of localization could be an artifact of this overexpression. Thus, although the genetic interactions indicate that EIPR-1 acts in the RAB-2 pathway, it is unclear whether EIPR-1 colocalizes with RAB-2 and its interactors.

### EIPR-1 interacts with members of the GARP and EARP trafficking complexes

WD40 domain proteins typically act as scaffolds that mediate multiple protein-protein interactions (28). Given the genetic interactions seen between *eipr-1* and members of the *rab-2* pathway, we asked whether the EIPR-1 protein shows physical interactions with any of the RAB-2 pathway proteins. Using yeast two-hybrid assays, we did not detect interactions between EIPR-1 and RAB-2, RUND-1 or CCCP-1. RAB-2 and RUND-1 were also localized normally in *eipr-1* mutants (data not shown). Thus, neither cellular localization nor physical interactions connect EIPR-1 directly to RAB-2 or its interactors, suggesting that EIPR-1 may act at a distinct step in this trafficking pathway.

To identify interactors of EIPR-1, we expressed GFP-tagged rat EIPR1 in the rat insulinoma cell line 832/13 and performed mass spectrometry on GFP pulldowns. 832/13 cells package insulin into dense-core secretory granules that closely resemble neuronal dense-core vesicles and share similar mechanisms of biogenesis, maturation, and release (29,30). Mammalian orthologs of RAB-2, RUND-1, CCCP-1 and EIPR-1 are all expressed endogenously in 832/13 cells (Figure 4A), so these cells are a reasonable place to look for interactors of these proteins.

**Figure 4.**
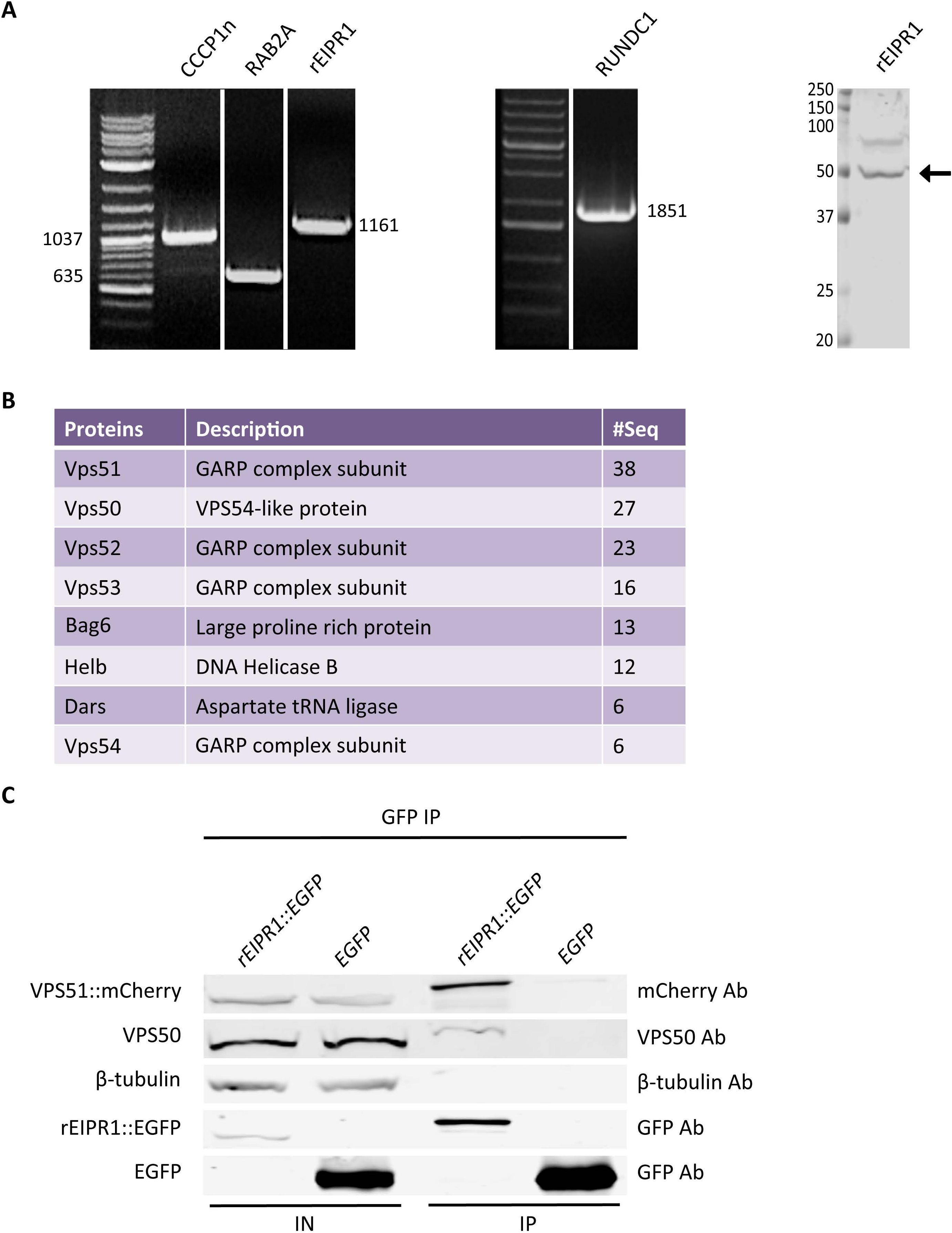
EIPR-1 interacts with members of the GARP and EARP complexes. (A) The rat ortholog of EIPR-1 is expressed in 832/13 cells. Left and middle: RT-PCR shows that rat orthologs of CCCP-1, RAB-2, EIPR-1 and RUND-1 are expressed in 832/13 cells. All bands are of the predicted size for the full-length cDNA except for CCCP1, which is truncated and corresponds to the N-terminal half of the gene. Right: Western blot shows expression of the rat EIPR-1 ortholog in 832/13 cells (arrow) at approximately its expected size (43 kD). (B) EIPR1 interacts with members of the GARP and EARP complexes. List of top hits from mass spectrometry of a pulldown of rEIPR1::GFP in 832/13 cells after subtracting hits found in GFP control pulldowns. # seq = number of unique peptides from each protein. All proteins with more than 5 unique peptides are shown from one of two independent experiments. More complete data tables for both experiments are shown in Figure S5. (C) EIPR1 interacts with VPS51 and VPS50. EGFP-tagged rat EIPR1 or EGFP was coexpressed with mCherry-tagged rat VPS51 in 832/13 cells. Immunoprecipitation of EIPR1::EGFP pulled down VPS51::mCherry and endogenous VPS50. Immunoprecipitation of untagged EGFP did not pull down VPS51::mCherry or VPS50. IN: input; IP: immunoprecipitation.

We performed two independent pulldowns of GFP-tagged rat EIPR1. The top four hits identify the proteins VPS51, VPS50 (also known as syndetin or VPS54L), VPS52, and VPS53 (Figure 4B; Figure S5). Also high on the list is VPS54. These five proteins define the related GARP and EARP trafficking complexes that share the VPS51, VPS52, and VPS53 proteins, but use either VPS54 or VPS50 as a fourth subunit (31–33). Thus, EIPR1 is likely tied closely to one or both of these complexes, neither of which has been previously linked to dense-core vesicle biogenesis. These proteomic results are especially striking because, as described below, we had independently identified the *vps-52* and *vps-53* genes in our forward genetic screen for dense-core vesicle mutants.

To confirm the interactions detected by mass spectrometry, we performed coimmunoprecipitation experiments. GFP-tagged EIPR1 expressed in 832/13 cells coimmunoprecipitated with either mCherry-tagged VPS51 or with endogenous VPS50 (Figure 4C), validating the interactions. Since no other proteins showed as strong an interaction by mass spectrometry as the GARP/EARP subunits, EIPR1 is a specific interactor of the GARP/EARP complexes.

### The GARP/EARP subunits VPS-51, VPS-52 and VPS-53 act in the RAB-2 pathway to control locomotion and dense-core vesicle cargo trafficking

Further supporting the model that EIPR-1 functions with either the GARP or EARP complex, we isolated mutants in the worm orthologs of *vps-52* and *vps-53* in our original forward genetic screen. We cloned these mutants by mapping, whole-genome sequencing, and transgenic rescue (see Materials and Methods). Both the *vps-52*(*ox345*) and *vps-53*(*ox339*) mutants are predicted to be molecular nulls, an early stop and deletion respectively, and both mutants phenocopy other deletion mutants in the same genes. These mutants also resemble a deletion mutant in the *vps-51* gene. Neuronal roles of *vps-51*, *vps-52*, and *vps-53* have not been previously described in *C. elegans*.

We tested whether *vps-51*, *vps-52* and *vps-53* mutants have behavioral and cellular phenotypes similar to those of *eipr-1* and *rab-2*. Like *eipr-1* and members of the *rab-2* pathway, *vps-51*, *vps-52* and *vps-53* mutants have slow but coordinated locomotion (Figure 5A) and an egg-laying defect. A double mutant between *rab-2* and *vps-53* did not have a stronger locomotion phenotype than the *rab-2* single mutant (Figure 5B), indicating that *rab-2* and *vps-53* act in the same pathway to control locomotion. These mutants also exhibit defects in trafficking dense-core vesicle cargos. Like *rab-2* and *eipr-1*, *vps-51*, *vps-52* and *vps-53* mutants all show reduced axonal levels of Venus-tagged NLP-21 or FLP-3 peptides (Figure 5C,D; Figure S6). A *rab-2*; *vps-53* double mutant did not show an enhanced NLP-21::Venus trafficking defect compared to the single mutants (Figure 5E), also suggesting that *rab-2* and *vps-53* act in the same pathway. However, unlike the *rab-2* and *eipr-1* mutants which show reduced axonal accumulation of the dense-core vesicle transmembrane protein IDA-1, mutants in *vps-51, vps-52* and *vps-53* all show increased axonal accumulation of IDA-1 (Figure 5F), indicating that these genes have at least some function distinct from *rab-2* or *eipr-1*. Since VPS-51, VPS-52, and VPS-53 are common to both GARP and EARP, these data suggest that GARP, EARP, or both are required for normal dense-core vesicle biogenesis and regulation of worm locomotion.

**Figure 5.**
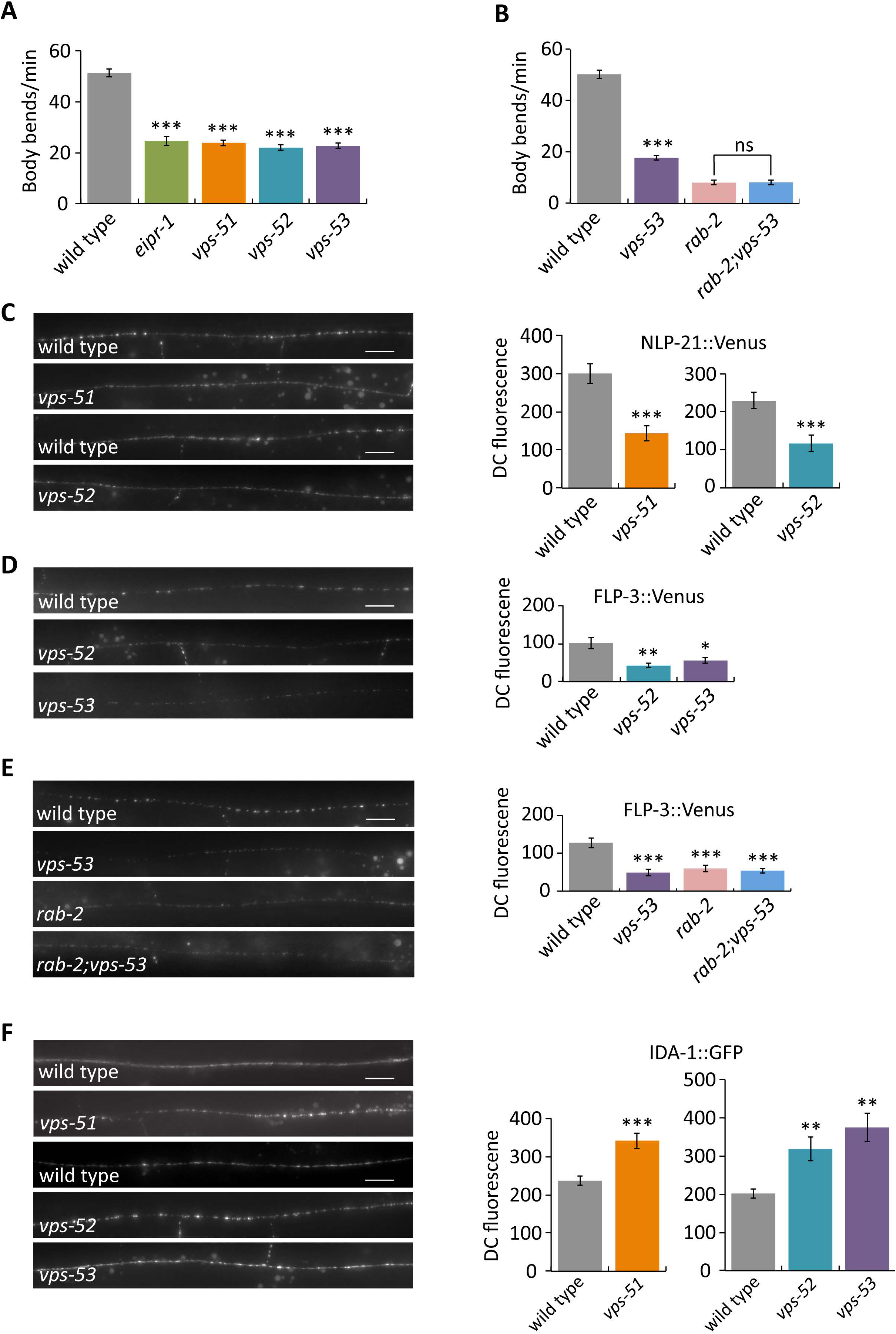
GARP/EARP mutants have defects in locomotion and trafficking dense-core vesicle cargos. (A) GARP/EARP mutants *vps-51*, *vps-52*, and *vps-53* have a reduced locomotion rate. ***, P<0.001 compared to wild type. Error bars = SEM; n = 10. (B) *vps-53* acts in the same genetic pathway as *rab-2* to control locomotion. A *rab-2; vps-53* double mutant does not have a stronger phenotype than a *rab-2* single mutant. ***, P<0.001 compared to wild type; ns, not significant, P>0.05. Error bars = SEM; n = 10. (C) *vps-51* and *vps-52* mutants have reduced levels of NLP-21::Venus fluorescence in the dorsal nerve cord. Left: representative images. Scale bar: 10 μm. Right: quantification. The mean dorsal cord (DC) fluorescence intensity is given in arbitrary units. *vps-51* and *vps-52* were assayed in separate experiments with independent matched wild type controls. ***, P<0.001 compared to wild type. Error bars = SEM; n = 9-10. (D) *vps-52* and *vps-53* mutants have reduced levels of FLP-3::Venus fluorescence in the dorsal nerve cord. Scale bar: 10 μm. ***, P<0.001; **, P<0.01 compared to wild type. Error bars = SEM; n = 10. (E) *vps-53* acts in the same genetic pathway as *rab-2* to control FLP-3::Venus trafficking. A *rab-2; vps-53* double mutant does not have a stronger phenotype than *rab-2* or *vps-53* single mutants. Scale bar: 10 μm. ***, P<0.001 compared to wild type. Error bars = SEM; n = 10. (F) *vps-51, vps-52* and *vps-53* mutants have increased levels of IDA-1::GFP in the dorsal cord. *vps-51* was assayed in a separate experiment from *vps-52* and *vps-53*, each with independent matched wild type controls. Scale bar: 10 μm. ***, P<0.001; *, P<0.05 compared to wild type. Error bars = SEM; n = 10.

To determine the expression pattern of *vps-52* and *vps-53*, we made transgenic worms expressing the YFP-derivative Citrine under either the *vps-52* or *vps-53* promoter. Both *vps-52* and *vps-53* were expressed strongly and widely in neurons and were also expressed in several other tissues (Figure 6A,B and Materials and Methods for details). These data are consistent with the reported expression patterns of *vps-51*, *vps-52*, and *vps-53* (34). To determine where VPS-52 and VPS-53 localize in neurons, we tagged them with tagRFP at their C-termini and expressed them under their own promoters as single-copy transgenes to achieve endogenous expression levels. Both VPS-52 and VPS-53 localized mainly to two or three perinuclear puncta in neuronal cell bodies (Figure 6C,D), reminiscent of the localization of RAB-2, RUND-1 and CCCP-1 to the trans-Golgi (23). We examined colocalization of VPS-52::tagRFP or VPS-53::tagRFP with GFP-tagged RAB-2 and found that they are partially overlapping in worm neurons (Figure 6C,D), consistent with the reported localization of GARP to the trans-Golgi in other cell types and other systems (Conibear and Stevens, 2000; Liewen et al., 2005; Lobstein et al., 2004; Luo et al., 2011).

**Figure 6.**
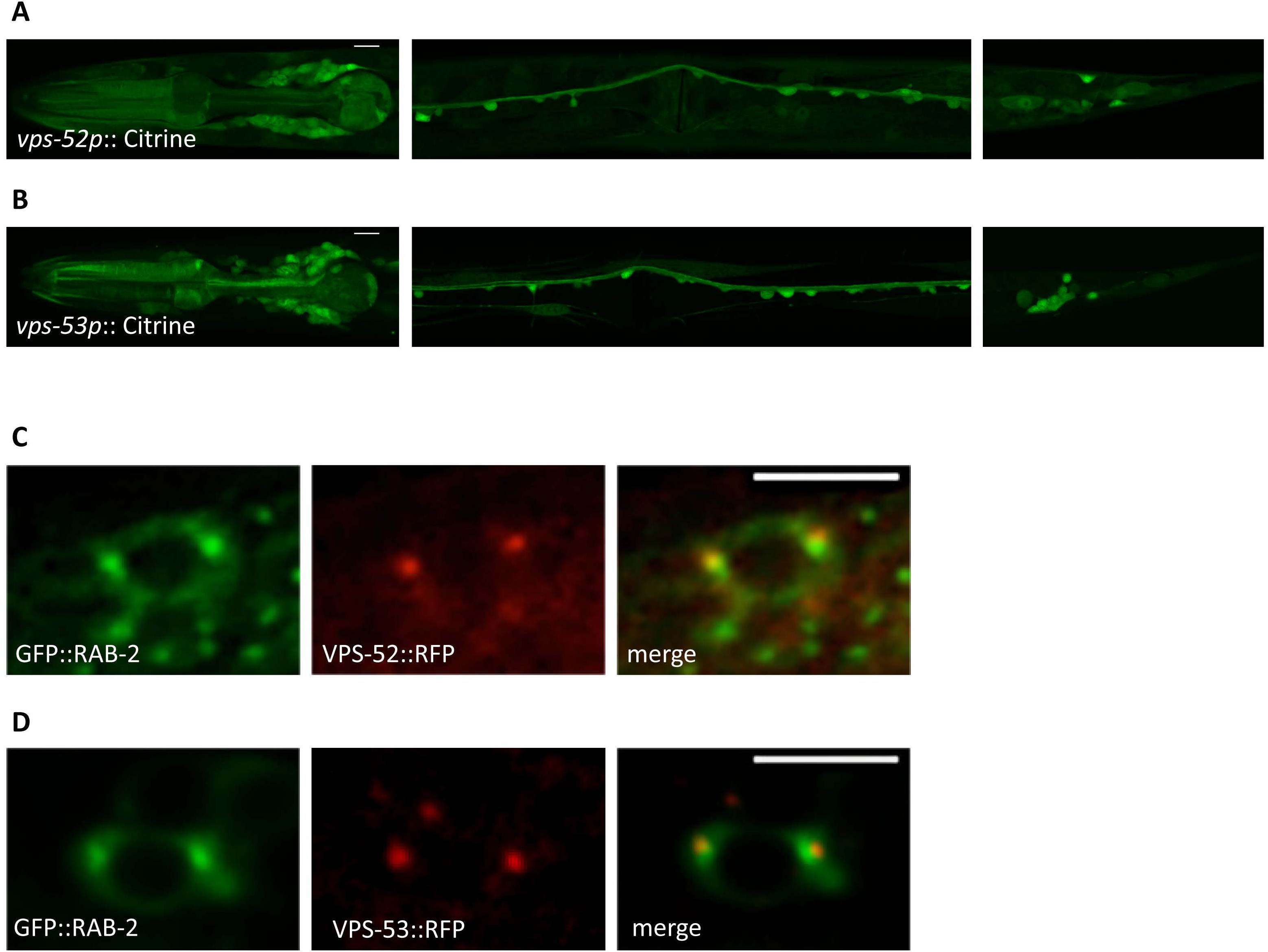
VPS-52 and VPS-53 are expressed in neurons and partially colocalize with RAB-2. (A) Representative images of animals expressing GFP under the *vps-52* promoter. The image on the left shows expression in the head neurons and pharynx, the image in the middle shows expression in ventral cord motor neurons and the image on the right shows expression in tail neurons. All images are single confocal slices of animals in a dorsal/ventral orientation. Scale bar: 10 μm. (B) Representative images of animals expressing GFP under the *vps-53* promoter. The image on the left shows expression in the head neurons and pharynx, the image in the middle shows expression in ventral cord motor neurons and the image on the right shows expression in tail neurons. All images are single confocal slices of animals in a dorsal/ventral orientation except the tail, which shows a lateral orientation. Scale bar: 10 μm. (C) Representative images of neurons coexpressing VPS-52::tagRFP and GFP::RAB-2. Scale bar: 5 μm. (D) Representative images of neurons coexpressing VPS-53::tagRFP and GFP::RAB-2. Scale bar: 5 μm.

### The EARP complex controls cargo sorting to dense-core vesicles

To determine whether it is GARP or EARP (or both) that is required for dense-core vesicle biogenesis, we obtained deletion mutations in the GARP-specific *vps-54* and EARP-specific *vps-50* subunits. Unlike the other GARP mutants, *vps-54* mutants did not have defects in locomotion or axonal accumulation of the luminal dense-core vesicle cargos NLP-21 and FLP-3 (Figure 7A,C; Figure S7A,B). In contrast, *vps-50* mutants had defects in locomotion and reduced axonal levels of luminal dense-core vesicle cargos very similar to those of mutants in *eipr-1*, *vps-51*, *vps-52* and *vps-53* (Figure 7A-C; Figure S7B). An *eipr-1*; *vps-50* double mutant did not have an enhanced defect in locomotion or dense-core vesicle cargo trafficking when compared to the *vps-50* single mutant (Figure 7A,B; Figure S7B), indicating that *eipr-1* and *vps-50* act in the same pathway. Moreover, like *eipr-1*, mutations in *vps-50* did not affect axonal levels of the synaptic vesicle protein synaptobrevin (Figure S7C), indicating that VPS-50 is not required for the trafficking of synaptic vesicle proteins. In 832/13 cells, endogenous VPS50 showed a perinuclear localization that partially overlapped GFP-tagged RAB2A and CCCP1 (Figure S8A,B). Providing additional evidence that *vps-50* acts in the dense-core vesicle pathway regulating worm locomotion, we found that a *vps-50* mutation suppressed the activated Gq mutant *egl-30*(*tg26*) that was used in the genetic screen for dense-core vesicle mutants. However, a *vps-54* mutation did not suppress this activated Gq mutant. Taken together, these data support a role for EARP, but not GARP, in dense-core vesicle biogenesis and regulation of worm locomotion.

**Figure 7.**
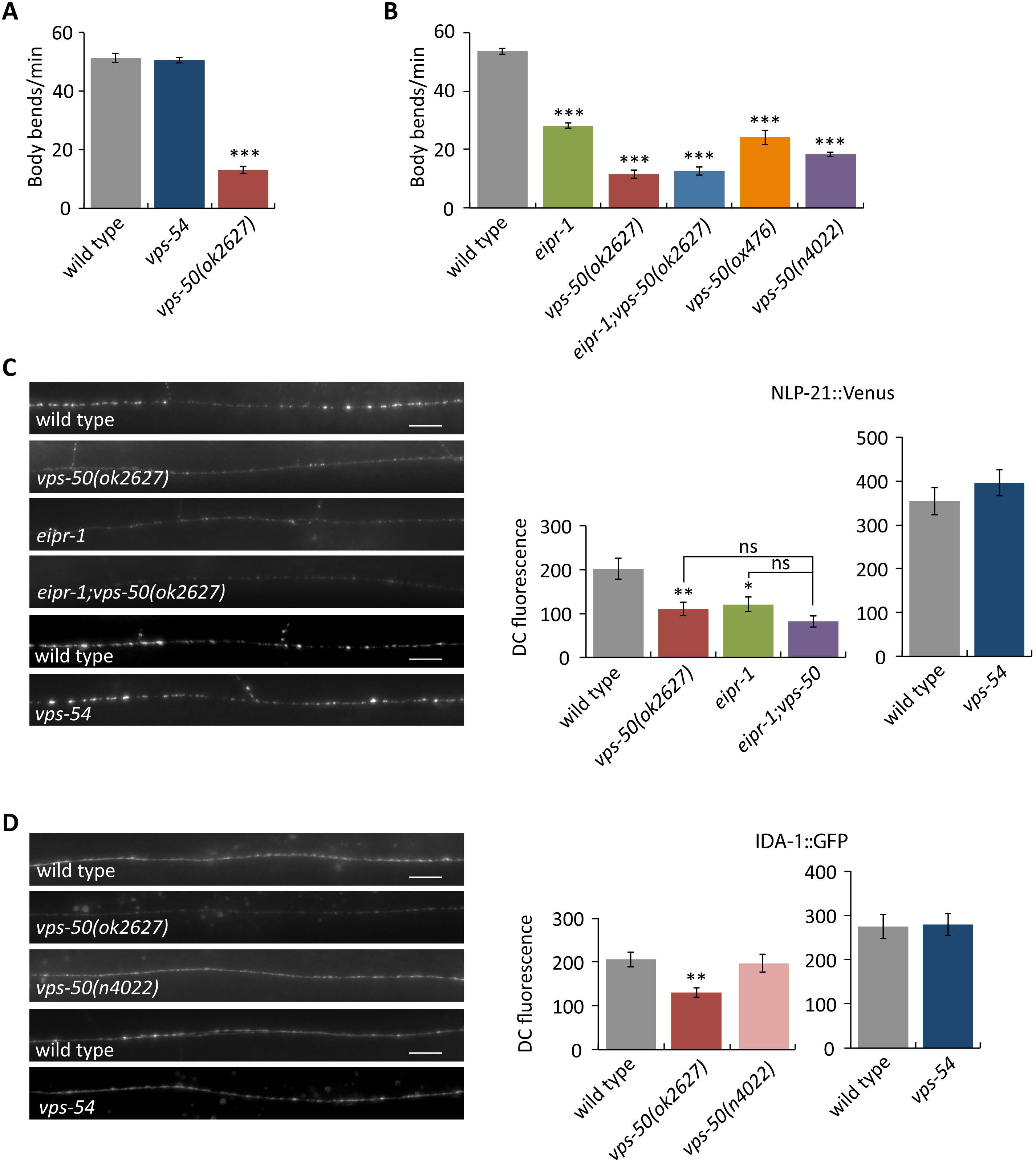
EARP, not GARP, is important for locomotion and trafficking dense-core vesicle cargos. (A) The EARP-specific mutant *vps-50* has a reduced locomotion rate, but the GARP-specific mutant *vps-54* does not have reduced locomotion. ***, P<0.001 compared to wild type. Error bars = SEM; n = 10. (B) *eipr-1* acts in the same genetic pathway as *vps-50* to control locomotion. An *eipr-1*; *vps-50* double mutant does not have a stronger phenotype than a *vps-50* single mutant. ***, P<0.001 compared to wild type. Error bars = SEM; n = 10. (C) The EARP-specific mutant *vps-50* has reduced levels of NLP-21::Venus fluorescence in the dorsal nerve cord, but the GARP-specific mutant *vps-54* does not. Left: representative images. Scale bar: 10 μm. Right: quantification. The mean dorsal cord (DC) fluorescence intensity is given in arbitrary units. *vps-50* and *eipr-1* have reduced NLP-21::Venus fluorescence, but an *eipr-1; vps-50* double mutant does not have a stronger phenotype than either single mutant (ns, not siginificant, P>0.05), indicating that *eipr-1* and *vps-50* act in the same genetic pathway. A *vps-54* mutant had no reduction in NLP-21::fluorescence. *vps-54* was assayed in a separate experiment with independent matched wild type control. **, P<0.01; *, P<0.05 compared to wild type. Error bars = SEM; n = 10-11. (D) IDA-1::GFP fluorescence levels in the dorsal cord. The *vps-50*(*ok2627*) deletion mutant had decreased IDA-1::GFP fluorescence (**, P<0.01) but the *vps-50*(*n4022*) late stop mutant and a *vps-54* deletion mutant had no decrease. *vps-54* was assayed in a separate experiment with independent matched wild type control. Scale bar: 10 μm. Error bars = SEM; n =10-12.

The decreased axonal levels of luminal dense-core vesicle cargos like NLP-21::Venus could be due either to decreased levels of the cargo in dense-core vesicles, or to increased release of the cargo. Exocytosis of dense-core vesicles and release of their cargos is reflected by accumulation of NLP-21::Venus in scavenger cells called coelomocytes that reside in the body cavity of the worm (35,36). Thus, we measured coelomocyte fluorescence as an indirect measure of dense-core vesicle release. Like *rab-2* mutants, *eipr-1*, *vps-50*, and *vps-52* mutants all showed reductions in the accumulation of Venus fluorescence in coelomocytes that is approximately proportional to the decrease in axonal fluorescence seen in these mutants (Figure S9). This suggests that these mutants are not defective in dense-core vesicle release per se, but release reduced amounts of dense-core vesicle cargo because the vesicles contain less cargo. Additionally, double mutants between *eipr-1* and *rab-2* or *vps-50* did not have stronger coelomocyte uptake phenotypes than the single mutants, providing further evidence that these genes all act in the same dense-core vesicle cargo sorting pathway.

Two independent *vps-50* mutants exhibited either a slight decrease or no decrease in axonal levels of the dense-core vesicle transmembrane cargo IDA-1 (Figure 7D). Thus, *vps-50* and *eipr-1* have a similar IDA-1 trafficking phenotype that is distinct from the increased axonal accumulation of IDA-1 seen in *vps-51*, *vps-52* and *vps-53* mutants. Interestingly, *vps-54* mutants showed no change in axonal accumulation of IDA-1 (Figure 7D).

Previously, *C. elegans* GARP mutants, including *vps-54*, were shown to have enlarged lysosomes in coelomocytes (34). We examined *eipr-1* and *vps-50* mutants for this phenotype using the lysosomal marker LMP-1::GFP expressed in coelomocytes. Unlike GARP mutants, neither *eipr-1* nor *vps-50* had enlarged lysosomes (Figure S10), indicating that EARP is not required for normal lysosomal morphology. Thus, GARP is required for normal lysosomal morphology and EARP is required for dense-core vesicle biogenesis.

## Discussion

In this study, we identified several new factors important for dense-core vesicle biogenesis: the WD40 protein EIPR-1 and the EARP trafficking complex with which it interacts. EIPR-1 and EARP are required for normal trafficking of dense-core vesicle cargos, and act in the same genetic pathway as the small GTP-binding protein RAB-2 and its interactors RUND-1 and CCCP-1. Loss of any of these proteins results in reduced levels of cargo in mature dense-core vesicles and associated behavioral defects. The identification of EARP, an endosomal recycling complex, suggests that dense-core vesicle cargo trafficking involves not only the forward trafficking of cargo into nascent dense-core vesicles as they bud off from the trans-Golgi, but also the retrieval or recycling of cargo or sorting factors from endosomal compartments (Figure 8).

**Figure 8.**
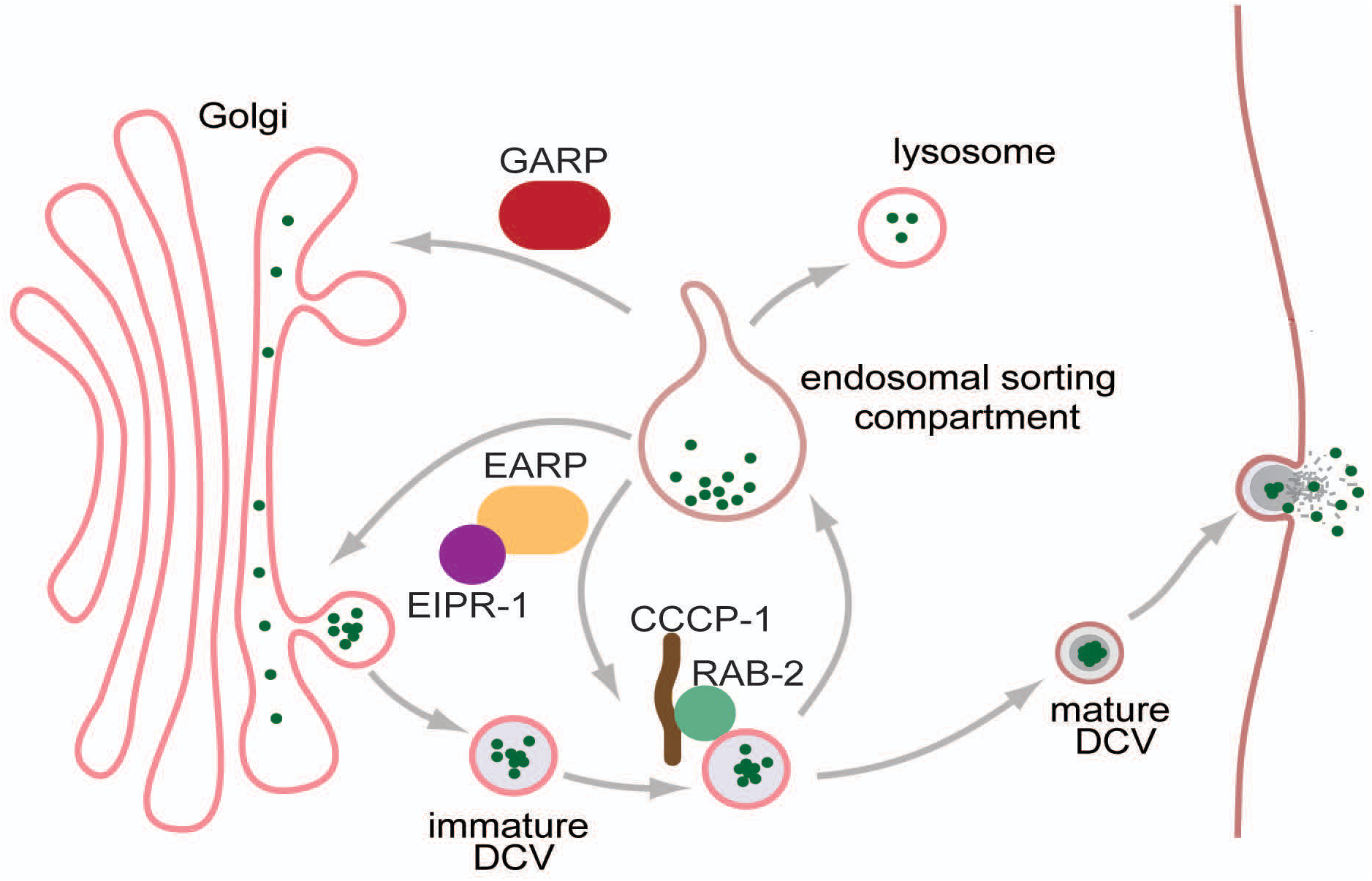
Model for dense-core vesicle cargo sorting by the EARP complex. This model combines and builds on aspects of the well-described ‘sorting by entry’ and ‘sorting by retention’ models for sorting cargos to dense-core vesicles (15,16). Aggregation of cargo (clustered green dots) within the trans-Golgi may initiate the process of cargo sorting (‘sorting by entry’). Then, following vesicle budding, non-aggregated cargo may be sorted away to endosomes while the aggregated core remains in the vesicle (‘sorting by retention’). Additional aggregation may occur in post-Golgi immature vesicles as the pH drops and propeptides are processed. Finally, EIPR-1/EARP and RAB-2 and its effectors (e.g. CCCP-1, shown) may mediate the retrieval of non-aggregated cargos from the endosomal compartment back to the trans-Golgi or immature DCVs. Such a cyclical sorting mechanism may serve both to remove non-DCV cargos and to provide non-aggregated DCV cargos a “second chance” at sorting to DCVs by recycling them for a new round of potential aggregation. Multiple rounds of such a cycle could lead to enriched retention of aggregated cargos in DCVs. In mutants of EARP or RAB-2, reduced retrieval of cargos from endosomes may lead to their ultimate loss to lysosomes and hence reduced cargo levels in mature DCVs. GARP acts in a distinct endosome to Golgi trafficking pathway. It is also possible that EARP and RAB-2 act by retrieving unidentified sorting receptors.

### The WD40 domain protein EIPR1 is a new interactor of the EARP complex

Through a convergence of worm genetics and proteomics in rat 832/13 cells, we identified the WD40 domain protein EIPR1 as a direct physical and functional interactor of the EARP complex. The EARP complex was recently identified as a new complex structurally related to the GARP complex (32,33). EARP shares the VPS51, VPS52 and VPS53 subunits with the GARP complex, but uses VPS50 instead of VPS54 as a fourth subunit. VPS50 was also recently shown to interact with GARP subunits in three large-scale mass spectrometry interactome data sets, demonstrating that the EARP complex is present in various cell types in several species (37–39). In two of these data sets, EIPR1 was shown to interact with EARP subunits in human HEK293T or HeLa cells (37,38), providing support for our results and indicating that the EIPR1 interaction with EARP is robust and conserved. In fact, using VPS50 as a bait protein, VPS51, VPS52, VPS53 and EIPR1 were pulled down as a stoichiometric complex (37), indicating that EIPR1 may form part of a stable complex with EARP. No VPS54 was pulled down. This is consistent with our genetic experiments showing that *eipr-1* and *vps-50* mutants have similar locomotion and dense-core vesicle cargo trafficking phenotypes that are shared by *vps-51*, *vps-52*, and *vps-53* mutants, but not *vps-54*. Thus, we propose that EIPR1 is a new member of the EARP complex.

The human *Eipr1* gene was originally named *Tssc1* for “tumor-suppressing subtransferable candidate 1” because it was first identified as one of several genes thought to be candidates for a tumor-suppressing activity on chromosome 11 (40). However, when the human genome was completed, *Tssc1* was located on chromosome 2, and thus could not be the putative tumor suppressor. Because the TSSC1 protein appears to be both physically and functionally connected to the EARP complex, we propose that it be renamed EIPR1 (EARP-Interacting PRotein).

WD40 domain proteins typically act as scaffolds that provide several interaction surfaces for the assembly of larger protein complexes (28). Though EIPR1 clearly interacts with EARP, it is unclear whether EIPR-1 is required for the localization or stability of the EARP complex. In an *eipr-1* mutant, VPS-52 still showed a punctate localization (Figure S11), though this could also reflect its participation in the GARP complex. In *C. elegans* (unlike yeast), the remaining members of the GARP complex were shown to be stable and localized in the absence of one of the subunits (34), so the same may hold true for EARP. Surprisingly, GFP-tagged EIPR-1 was found throughout the cytoplasm when overexpressed in *C. elegans* neurons and not associated with a membrane-bound compartment. However, overexpression of membrane-localized proteins can sometimes lead to a diffuse cytoplasmic appearance (41). Because we could not detect an endogenously tagged EIPR-1::GFP fusion protein, we presume that *eipr-1* is normally expressed at low levels. This conclusion is supported by the few cDNAs or RNAseq reads reported for *eipr-1* on Wormbase (http://www.wormbase.org/species/c_elegans/gene/WBGene00013599#0df9-gec6-2).

One other hit found in both of our EIPR1 pulldowns is SNAP29, a late Golgi/endosomal SNARE protein that has been shown to function in endocytic recycling and autophagosome fusion (42–45). SNAP29 was also identified as an interactor of EIPR1 or EARP subunits (but not VPS54) in other recent mass spectrometry studies (33,37,38), and we confirmed that SNAP29 interacts with EIPR1 by a coimmunoprecipitation experiment (Figure S12). GARP has been shown to interact with several late-Golgi SNARE proteins and has been proposed to both tether vesicles and promote SNARE assembly that leads to fusion (34,46–49). We propose that SNAP29 is part of a SNARE complex specifically interacting with EARP.

### A new role for the GARP/EARP complex in mediating dense-core vesicle cargo sorting

EARP was originally shown to be localized to a Rab4 positive endosomal compartment in both fly S2 cells and rat hippocampal neurons (32,33), and depletion of the EARP-specific VPS50 subunit led to a modest defect in the endocytic recycling of the transferrin receptor back to the cell surface (33). Here we identify a new neuronal role of this complex in the sorting of cargos to dense-core vesicles.

The GARP complex was originally characterized as an endosome to Golgi retrograde trafficking complex that may tether endosome-derived vesicles at the trans-Golgi (46,48,50,51). Loss or depletion of GARP subunits leads to a variety of phenotypes that are presumed to be secondary to a primary defect in retrograde trafficking, including missorting of lysosomal proteins, abnormal lysosomal morphology, defective autophagy, defective lipid and sterol transport, defective anterograde transport of glycosyl phosphatidylinositol (GPI)-anchored and transmembrane proteins, and defects in sphingolipid homeostasis (31,34,47,50–53).

GARP and EARP subunits are expressed especially strongly in the nervous system in both mice and worms (33,34,54,55), but until now, neuronal phenotypes of GARP/EARP mutants have not been carefully characterized. However, several mutations in GARP/EARP subunits have been connected to neurological diseases. Mutations in VPS54 and VPS52 lead to neurological disorders in mice that have been studied as models of the human diseases amyotrophic lateral sclerosis (ALS) and the seizure disorder high-pressure nervous syndrome, respectively (56,57). Also, mutations in the human VPS53 gene cause progressive cerebello-cerebral atrophy type 2 (PCCA2), a neurodegenerative disease characterized by severe mental retardation and early-onset epilepsy (58). It will be interesting to determine whether the GARP/EARP mutations cause these diseases due to their previously characterized role in sorting lysosomal cargos (50,51), or due to their newly identified role in dense-core vesicle biogenesis. It will also be interesting to determine whether neurological diseases associated with VPS52 and VPS53 are caused by dysfunction of GARP or EARP or both. The overt behavioral phenotypes of *C. elegans* mutants affecting the GARP/EARP common subunits (*vps-51*, *vps-52* and *vps-53*) are very similar to those of *vps-50* and *eipr-1*, suggesting that EARP is more important than GARP for neuronal function in *C. elegans*.

Our identification of EIPR-1 and EARP as important for dense-core vesicle cargo sorting supports a recent study that showed that *vps-50* mutants have locomotion and dense-core vesicle cargo trafficking defects (55). Like *eipr-1*, the slow locomotion of *vps-50* mutants on food was partially rescued by expression in cholinergic neurons (55), suggesting that *eipr-1* and *vps-50* function together in the same neurons to control locomotion behavior. VPS-50 was also shown to physically interact with the vacuolar-ATPase (v-ATPase) complex that acidifies dense-core vesicles, and *vps-50* mutants had defects in vesicle acidification and assembly of the v-ATPase complex (55). These defects may be indirectly caused by a primary defect of EARP mutants in missorting dense-core vesicle cargos, possibly including subunits of the v-ATPase.

How does EARP sort dense-core vesicle cargos? Thus far, all of the factors in the RAB-2/EARP pathway are cytoplasmic proteins that localize to membranes but do not have transmembrane domains and thus cannot communicate directly with luminal cargos inside the vesicles. Thus, either there is a transmembrane sorting receptor yet to be identified or the RAB-2/EARP pathway proteins help sort cargo by a passive mechanism that does not involve direct interactions with cargo. We favor a model based on the latter idea (Figure 8). This model combines and builds on aspects of the well-described ‘sorting by entry’ and ‘sorting by retention’ models for cargo sorting to dense-core vesicles (15,16). Specifically, we propose that cargo sorting occurs through the following steps: (1) an intrinsic ability of certain dense-core vesicle cargos to aggregate in the trans-Golgi (sorting by entry), (2) additional aggregation in immature dense-core vesicles as the luminal pH is reduced and propeptides are processed into their mature forms, and (3) the repeated recycling of non-aggregated cargo from immature dense-core vesicles through an endosomal compartment back to the trans-Golgi/immature dense-core vesicles. Such a recycling pathway will help sort away soluble non-dense-core vesicle cargos (sorting by retention), and also help concentrate aggregated dense-core vesicle cargos by providing these cargos multiple chances to aggregate and be retained by mature dense-core vesicles. We propose that EARP and the RAB-2 pathway may control the trafficking into or out of the endosomal compartment. In mutants in this pathway, increased trafficking of cargos to endosomes or reduced retrieval of cargos from endosomes leads to their ultimate loss to lysosomes and hence reduced cargo levels in mature dense-core vesicles. Thus, EARP and the RAB-2 pathway do not act instructively to sort cargos in the way a sorting receptor would, but instead have a permissive role, somewhat analogous to the mechanism by which chaperones assist protein folding by providing proteins multiple chances to fold correctly and avoid degradation. Such a permissive role for EARP and the RAB-2 pathway would be consistent with the fact that none of these mutants has a complete loss of cargo sorted to the mature dense-core vesicles that accumulate in axons.

Several previous studies provide support for our model. Following dense-core vesicle exocytosis, the dense-core vesicle transmembrane protein peptidylglycine a-amidating monooxygenase (PAM) has been shown to be recycled back to dense-core vesicles via endosomal compartments (59), demonstrating that there must be trafficking of cargos from endosomes to dense-core vesicles. Our model builds on this observation to suggest that such endosomal trafficking occurs not only following exocytosis, but also during the biogenesis of dense-core vesicles. Another study demonstrated that soluble dense-core vesicle cargos are segregated from self-aggregating cargos in immature dense-core vesicles near the trans-Golgi (60), consistent with the idea that aggregation of cargos at this point in vesicle maturation can be sufficient to drive sorting into mature dense-core vesicles.

### A growing number of proteins important for dense-core vesicle biogenesis

The identification of EIPR-1 and EARP adds to a growing list of proteins that have been recently identified as being involved in mediating aspects of dense-core vesicle biogenesis. In *C. elegans*, we and others previously identified the small GTPase RAB-2; the putative RAB-2 GAP TBC-8; the RAB-2 effectors RIC-19, RUND-1 and CCCP-1; and the membrane-associated Golgi-localized protein HID-1 (19,20,22,23,61,62). Mutations in all of these proteins result in similar defects in dense-core vesicle cargo trafficking, and genetic epistasis data support the idea that they function in a common pathway.

In other systems, a number of factors have been recently identified as having roles in dense-core vesicle biogenesis. These include factors that belong to protein families implicated in various aspects of membrane trafficking such as BAR domain proteins (18,21,63,64), members of the Arf family of small GTPases (65), the clathrin adaptor complexes AP-1 and AP-3 (66–69), members of the BLOC-1 and HOPS complexes that are important for the biogenesis of lysosome-related organelles (70–72), and members of the sortilin family of lysosomal sorting receptors (73,74). Among these, there are a few possible connections to the RAB-2/EARP pathway we have described in *C. elegans.* AP-1 is involved in trafficking between various cellular compartments, most notably between the trans-Golgi and endosomes (75), and has been shown to be associated with immature dense-core vesicles (6,8), so it may be involved in the endosomal recycling pathway proposed by our model. The BLOC-1 subunit dysbindin was identified as an interactor of EIPR1, VPS51 and VPS53 by mass spectrometry (38), and the possible dense-core vesicle coat protein VPS41 (along with other HOPS subunits) was identified as an interactor of Rab2 (32).

One final factor shown to be important for dense-core vesicle biogenesis in mouse chromaffin cells is the SNARE protein Vti1a (76). Interestingly, Vti1a is one of the components of a trans-Golgi SNARE complex that binds GARP (48), and it also localizes to immature dense-core vesicles near the trans-Golgi (76). Syntaxin 6, another member of this SNARE complex, has also been shown to be associated with immature dense-core vesicles and small vesicles near the trans-Golgior endosomes (8,11,77). Blocking antibodies to syntaxin 6 were shown to inhibit the homotypic fusion of immature dense-core vesicles in an *in vitro* fusion assay (11), but the physiological importance of syntaxin 6 for dense-core vesicle biogenesis *in vivo* remains unclear. Interestingly, the RAB-2 effector RUND-1 is tightly colocalized with syntaxin 6 in *C. elegans*neurons (23) and the BAR domain protein PICK1 tightly colocalized with syntaxin 6 in mouse pituitary cells (21). Also, syntaxin 6 physically interacts with SNAP29 (11), a SNARE protein that we and others identified as an EIPR1 or EARP interactor (33,37,38). It will be interesting to determine whether SNAP29, Vti1a, and syntaxin 6 are members of a SNARE complex that interacts with EARP, and the compartment where they function.

## Materials and methods

### Strains

Worm strains were cultured and maintained using standard methods (Brenner, 1974). A complete list of strains and mutations used is provided in the Strain List (Table S1).

### Isolation and identification of *eipr-1*, *vps-52*, and *vps-53* mutations

The *eipr-1*(*ox316*), *vps-52*(*ox345*) and *vps-53*(*ox339*) mutants were isolated in an ENU screen for suppressors of the hyperactive locomotion of an activated Gq mutant, *egl-30*(*tg26*) (23). When outcrossed away from the *egl-30*(*tg26*) mutation, all three of these mutants had sluggish unmotivated locomotion and defects in egg-laying. The phenotypes of all three of these mutants are partially maternally rescued; for example, the phenotype of an *eipr-1* mutant is weaker if its parent is heterozygous (*eipr-1/+*) than if its parent is homozygous (*eipr-1/eipr-1*). The basis of this maternal rescue is not clear, but indicates the likely maternal deposition of these proteins in the egg.

We mapped these mutations using single nucleotide polymorphisms (SNPs) in the Hawaiian strain CB4856 as described (Davis et al., 2005). The *eipr-1*(*ox316*) mutation was mapped to a 205 kb region on the right arm of chromosome I with 31 predicted protein-coding genes. Because there was no coverage of this region by *C. elegans* cosmids, we obtained *C. briggsae* BACs (from CHORI) carrying orthologs of the genes in this region and injected them into *eipr-1(ox316)* mutants. The BAC RPCI94_01L06 rescued the *eipr-1* locomotion defect, while the BAC RPCI94_13K10 did not. RPCI94_01L06 carries orthologs of six *C. elegans* genes in the mapped region. We sequenced the best two candidate genes in the *ox316* mutant and found a C to T transition mutation in exon 4 of Y87G2A. 11, leading to a premature stop codon at Q310. We obtained and sequenced the *eipr-1* cDNA from the ORFeome library, confirming the gene structure predicted on Wormbase. The clone carried a single mutation that we fixed by Quick Change mutagenesis. We then cloned the mutation-free cDNA into a Gateway entry vector for use in rescue experiments. We rescued the *ox316* mutant with a transgene carrying only Y87G2A. 11, confirming the gene identification.

The *vps-52*(*ox345*) mutation was mapped to a 720 kb region in the middle of the X chromosome with 122 genes. The *vps-53(ox339)* mutation was mapped to a 92 kb region in the middle of chromosome III with 23 genes. We constructed a double mutant strain carrying both the *ox339* and *ox345* mutations and performed Illumina whole-genome sequencing. Within the region where *ox339* mapped, we found no SNPs predicted to alter protein function, but did identify a 687 bp deletion within the T05G5.8 (*vps-53*) ORF. Within the region where *ox345* mapped, we found only a single SNP predicted to alter protein function, a C to T transition mutation in the F08C6.3 (*vps-52*) ORF that created a premature stop codon at S119. The mutations in the *vps-53* and *vps-52* genes were confirmed by Sanger sequencing. We confirmed the gene identifications by performing complementation tests with other deletion mutations in each gene and by performing single-gene rescue experiments.

### Molecular biology and transgenes

A complete list of constructs is provided in the Plasmid List (Table S2). Most of the constructs were made using the three slot multisite Gateway system (Invitrogen). Typically, a promoter, a coding sequence (genomic DNA or cDNA), and an N‐ or C-terminal fluorescent tag (GFP or tagRFP-T) were cloned along with a 3’UTR into the pCFJ150 destination vector used for Mos1-mediated single copy insertion (MosSCI) on chromosome II at *ttTi5605* (78,79). All insertions were made by the direct injection MosSCI method. For most constructs, we isolated multiple independent insertions that behaved similarly. Extrachromosomal arrays were made by standard transformation methods (80).

### Expression of *eipr-1*, *vps-52*, and *vps-53*

*eipr-1* is the fourth gene in an operon about 30 kb downstream from the start of the first gene in the operon, so it is not clear what promoter regulates *eipr-1.* We made a construct driving expression of GFP by the promoter region upstream of the first gene of the operon (1348 bp upstream of the W09C5.1 start codon). Extrachromosomal arrays of this construct showed expression in several tissues including the intestine, pharynx and hypodermis (Figure S4A), but not neurons, suggesting that this promoter fusion does not represent the endogenous expression pattern of *eipr-1*. We also tagged the chromosomal *eipr-1* gene with GFP using the CRISPR/Cas9 method (81). However, neither N-terminal nor C-terminal GFP-tagged *eipr-1* was bright enough to detect, even with the use of an anti-GFP antibody to amplify the signal, indicating that *eipr-1* is likely to be expressed endogenously at low levels.

We made constructs driving expression of the yellow-fluorescent protein Citrine under the *vps-52* and *vps-53* promoters and generated worms with extrachromosomal arrays. For *vps-52*, 822 bp upstream of the start codon was used as a promoter. *vps-53* is the second gene in an operon downstream of the gene *gcc-2*. For *vps-53*, 4502 bp upstream of the *gcc-2* start codon was used as the promoter. *vps-52* showed strong expression in neurons in the head, ventral cord, and tail (Figure 6A), as well as expression in the pharynx, intestine, uterus, spermatheca, skin, and several muscle types (body wall, vulval). Other than the skin and muscle expression, this expression pattern matches that of *rund-1* (23). *vps-53* was not as strongly expressed, but was clearly visible in many head neurons (though possibly not all), ventral cord neurons, tail neurons, the pharynx, and weakly in the body wall muscles (Figure 6B).

### Locomotion and egg-laying assays

To measure locomotion, first-day adults were picked to thin lawns of OP50 bacteria, allowed to rest for 30 seconds, and then body bends were counted for one minute. A body bend was defined as the movement of the worm from maximum to minimum amplitude of the sine wave (82).

To measure egg-laying, L4 larvae were placed on plates with OP50 and allowed to mature at 25°C overnight. The next day, five adult animals were moved to a fresh plate and allowed to lay eggs at 25°C for 2.5 hours. The number of eggs present on the plate was then counted.

### Cell culture

The INS-1-derived 832/13 cell line (83) was obtained from Dr. Christopher Newgard (Duke University School of Medicine) via Dr. Ian Sweet and Dr. Duk-Su Koh (University of Washington). 832/13 cells were routinely grown at 5% CO2 at 37°C in RPMI-1640, GlutaMAX^™^ (GIBCO), supplemented with 10% FBS, 1 mM Sodium pyruvate, 10 mM HEPES, 0.0005% 2-beta-mercaptoethanol, and 1X Pen/Strep (GIBCO). Cells were passaged biweekly after trypsin-EDTA detachment. All studies were performed on 832/13 passages between 70 and 90. 832/13 cells were transfected using Lipofectamine 2000 (ThermoFisher) according to the manufacturer’s instructions.

### Expression and purification of anti-GFP nanobody

The anti-GFP nanobody was expressed and purified as previously described (84) with few modifications. In brief, bacterial expression vector pLaG16 containing the anti-GFP nanobody was transformed in Arctic Express (DE3) cells (Agilent). Cells were induced with 0.1 mM IPTG for 16 h at 8°C and centrifuged at 5,000g for 10-min at 4°C. To isolate the periplasmic fraction by osmotic shock (85), cells were incubated for 1h in TES buffer (0.2 M Tris-HCl pH 8, 0.2 mM EDTA and 0.5 M sucrose). Cells were spun down for 10 min at 6,000g at 4°C and the pellet was suspended in TE buffer (0.2 M Tris-HCl pH 8 and 0.5 mM EDTA) following a 45 min incubation. Cells were pelleted by a 30 minutes spin at 30,000g at 4°C. The supernatant (periplasmic fraction) was bound to PerfectPro Ni-NTA Agarose affinity resin (5 Prime) for 1 h at 4°C. The resin was washed once with wash buffer I (20 mM sodium phosphate pH 8.0, 0.9 M NaCl) and twice with wash buffer II (20 mM sodium phosphate pH 8.0, 150 mM NaCl and 10 mM imidazole) and eluted with His elution buffer (20 mM sodium phosphate pH 8.0, 150 mM NaCl, 250 mM imidazole). The eluent was then dialyzed with PBS. Recombinant anti-GFP nanobody was conjugated to epoxy-activated magnetic Dynabeads M270 (Life Technologies) using 10 μg recombinant protein per 1 mg of Dynabeads, with conjugations carried out in 0.1 M sodium phosphate pH 8.0 and 1 M ammonium sulfate, with an 18 h incubation at 30°C. Beads were washed once with 100 mM Glycine pH 2.5, once with 10 mM Tris pH 8.8, 4 times with PBS and twice with PBS plus 0.5% Triton X-100. Beads were stored at −20°C in PBS with 50% glycerol.

### Mass spectrometry

For immunoprecipitation, 4×10^6^ 832/13 cells were plated onto 10 cm petri dishes. Twenty-four hours later, cells were transfected with either EIPR1::GFP or GFP. After 24 hours, cells were washed with PBS and harvested in lysis buffer [50 mM Tris pH 7.5, 150 mM NaCl, 1% NP-40, Protease inhibitor cocktail (Pierce)]. Lysates were passed 10 times through a 20G needle and incubated for 30 min at 4°C. Lysates were pre-cleared by a 20,000 g centrifugation at 4°C for 15 minutes and the supernatant was incubated with 30 μg of anti-GFP nanobody bound to magnetic beads, for two hours at 4°C. The beads were washed three times with 250 μL wash buffer (50 mM Tris pH 7.5, 300 mM NaCl, 1% NP-40). Samples were eluted in two steps with (1) 0.2 M glycine pH 2.5 and (2) 1 M NH4OH. Eluents were neutralized with 0.33 M Tris-HCl pH 7.5 and stored at - 80°C.

Samples were processed using modified FASP (filter aided sample preparation) to remove mass spec incompatible substances (86). Samples were filtered with 10K Amicon Ultra 0.5 ml filters (Millipore). Samples were buffer exchanged with 8 M urea in 50 mM ammonium bicarbonate pH 7.8, reduced with TCEP (Pierce), alkylated with IAA (Sigma) and digested with trypsin (Pierce). Samples were concentrated by speed vacuum.

A total of 0.15 μg of peptides was chromatographically separated onto a Thermo LTQ-Velos Pro mass spectrometer coupled with a Waters nanoACQUITY liquid chromatography system. A 75 μm fused silica column (Polymicro Technologies) was loaded with 30 cm C12 Jupiter (Phenomenex), 4 μm reverse-phase beads, and a 100 μm fused silica Kasil frit trap (PQ Corporation) loaded with 4 cm of C12 Jupiter. Peptides were separated by a 110 minute gradient consisting of buffer A (95% water, 5% acetonitrile and 0.1% formic acid) and buffer B (95% acetonitrile, 5% water and 0.1% formic acid). Two analytical replicates were run for each sample with each set of replicates randomized. Quality control samples were run every sixth sample and at the beginning of the runs to assess column chromatography stability.

Data were searched with SEQUEST (87) against a *Rattus norvegicus* FASTA database containing contaminants. False discovery rates were determined via a decoy database using Percolator (88) at a q-value threshold of 0.01 and peptides were assembled into protein identifications using ID Picker (89). Lists of EIPR1::GFP hits were assembled after first subtracting hits found in a GFP control pulldown (Figure S5A) or subtracting hits found in the pulldowns of two other proteins, CCCP1::GFP and RUNDC1::GFP (Figure S5B), performed in parallel.

### Coimmunoprecipitation and immunoblotting

For co-immunoprecipitation, 832/13 cells were cultured and lysed as described above (Mass spectrometry). The lysates were incubated with 20 μg anti-GFP nanobody bound to magnetic beads, the beads were washed three times with lysis buffer and resuspended in Laemmli loading buffer. Samples were resolved on 10% SDS-polyacrylamide gels and blotted onto nitrocellulose or PVDF membranes. To detect co-precipitated proteins, we added the following primary antibodies: mouse monoclonal anti-GFP (1:1000, Santa Cruz #sc-9996), monoclonal anti-mCherry (1:50, a gift from Jihong Bai and the Fred Hutchinson Cancer Research Center antibody development shared resource center), monoclonal anti-VPS50 (1:1000, FLJ20097 monoclonal antibody, clone 2D11, Abnova #H00055610-M01), rabbit polyclonal anti-SNAP29 (1: 1000, Sigma #S2069), or monoclonal anti-beta-tubulin as a loading control (1:1000, ThermoFisher, BT7R, #MA5-16308). The secondary antibody was an Alexa Fluor 680-conjugated goat anti-mouse antibody (1:20,000, Jackson Laboratory #115-625-166) or Alexa Fluor 680-conjugated goat anti-rabbit antibody (1:20,000, Jackson Laboratory #115-625-144). A LI-COR processor was used to develop images.

### Imaging and image analysis

Worms were mounted on 2% agarose pads and anesthetized with 50 mM sodium azide. Images were obtained using a Nikon 80i wide-field compound microscope except for the images shown in Fig 6A&B which were obtained on an Olympus confocal microscope, and the images in Fig 6C&D which were obtained on a Deltavision deconvolution microscope. To image the dorsal nerve cords, young adult animals were oriented with dorsal side up by exposure to the anesthetic for ten minutes on the slide before placing the cover slip. For quantitative imaging of dorsal cord fluorescence, all strains in a given experiment were imaged on the same days and all microscope settings were kept constant. The same section of the dorsal cord around the vulva was imaged in all worms. Maximum intensity projections were quantified using ImageJ software, measuring the total fluorescence in a region of interest encapsulating the cord and subtracting the background fluorescence of a region of identical size adjacent to the cord.

### Immunostaining of 832/13 cells

2×10^5^ 832/13 cells per well were seeded onto sterilized cover slips placed in 12 well cell culture plates. The next day, cells were rinsed twice with ice-cold PBS and fixed with 4% paraformaldehyde in PBS for 20 minutes at room temperature. The cells were rinsed twice with PBS and permeabilized with 0.5% Triton X-100 in PBS for 5 minutes at room temperature. The cells were again washed twice with PBS and then placed in 5% milk in PBS for 1 hour at room temperature, followed by immunofluorescence staining with rabbit polyclonal anti-CCDC132 (1:50, Sigma #HPA026679) and mouse monoclonal anti-GFP (1:350, Santa Cruz #sc-9996) in 0.5% milk in PBS at room temperature for 1 hour. The cells were then washed with PBS three times for 5 minutes each, and incubated with Rhodamine anti-rabbit secondary antibody (1:1000, Jackson Immunoresearch #111-025-144) and Alexa Fluor 488 anti-mouse secondary antibody (1:1000, Jackson Immunoresearch #115-545-146) at room temperature for 1 hour. The cells were washed with PBS three times for 5 min each and examined by fluorescence microscopy (Nikon 80i wide-field compound microscope). The anti-CCDC132 antibody recognizes VPS50.

### Statistics

P values were determined using GraphPad Prism 5.0d (GraphPad Software). Data sets with multiple comparisons were analyzed by a one-way ANOVA followed by a Bonferroni posthoc test to examine selected comparisons or by Dunnett’s test if all comparisons were to the wild type control. Pairwise data comparisons were analyzed by two-tailed unpaired t tests.

## Acknowledgments

We thank ShoheiMitani for the *eipr-1*(*tm4790*) mutant; Nicolas Paquin and Bob Horvitz for the *vps-50*(*n4022*) mutant; Christopher Newgard, Ian Sweet and Duk-Su Koh for the 832/13 cell line, with the support of the UW DRC Cell Function and Analysis Core (DK17047); King Yabut and Trisha Davis for the GFP nanobody and advice on mass spectrometry; JihongBai and the Fred Hutch Antibody Technology Resource for the anti-mCherry antibody; Suzanne Hoppins for the pmCherry-N1 vector; Colin Thacker and Richard Clark for assistance with Illumina sequencing; JihongBai, Gunther Hollopeter, and Alex Merz for comments on the manuscript; and Erik Jorgensen, in whose lab early phases of this project were performed. Some strains were provided by the CGC, which is funded by NIH Office of Research Infrastructure Programs (P40 0D010440).

## Supporting information captions

**S1 Figure. *eipr-1* mutants have egg laying defects.**

(A) *eipr-1* acts in neurons to control egg laying. The graph shows the number of eggs laid by 5 animals in a 2.5 hour period. *eipr-1* mutants show egg-laying defects and this phenotype is rescued by panneuronal expression of either the worm gene or its mouse ortholog (***, P<0.001). Error bars = SEM; n =10 plates of 5 worms each.

(B) *eipr-1* acts in the same pathway as *rab-2* to control egg-laying. Double mutants of *eipr-1* with *rab-2* or *rund-1* do not have stronger egg-laying defects than the single mutants. Though *rund-1* mutants have a visible Egl (egg-laying defective) phenotype indicating that they retain more eggs, they did not have a significantly reduced egg-laying rate as measured by this assay. Error bars = SEM; n = 5 plates of 5 worms each. ns, not significant, P>0.05.

**S2 Figure. Alignment of EIPR-1**. Alignment of *C. elegans* EIPR-1 (worm, accession # NP_493383.1) and its orthologs from *Arabidopsis thaliana* (Arabidopsis, accession # NP_173478.2), *Drosophila melanogaster* (fly, CG10646, accession # NP_648581.1), and *Homo sapiens* (human, TSSC1, accession # NP_003301.1). Identical residues are shaded in black and similar residues are shaded in gray. The WD40 repeats (from SMART, using the worm protein; http://smart.embl-heidelberg.de/) are marked with single black bars. Using SMART, worm EIPR-1 has four predicted WD40 repeats, Arabidopsis has six, fly has three, and human has five. WD40 repeats are difficult to identify by primary sequence and are often missed by prediction programs. The position of the ox316 stop mutation is marked with an asterisk. Alignment was made with MUSCLE (http://www.ebi.ac.uk/Tools/msa/muscle/) using default parameters and exhibited with Boxshade 3.21 (http://www.ch.embnet.org/software/BOX_form.html).

**S3 Figure. *eipr-1* mutants have defects in trafficking dense-core vesicle cargos but not synaptic vesicle cargos.**

(A) Representative images of FLP-3::Venus fluorescence in dorsal cord motor neuron axons of the wild type and *eipr-1*(*tm4790*) mutant strains. Scale bar: 10 μm. *eipr-1* mutants have decreased fluorescence in the dorsal cord, indicative of a FLP-3::Venus sorting or trafficking defect.

(B) Representative images of INS-22::Venus fluorescence in dorsal cord motor neuron axons. Scale bar: 10 μm.

(C) Representative images of IDA-1::GFP fluorescence in the dorsal nerve cord. Scale bar: 10 μm.

(D) SNB-1::GFP fluorescence levels in the dorsal nerve cord. *eipr-1* mutants do not have a defect (ns = not significant, P>0.05). Error bars = SEM; n =21-26.

**S4 Figure. Expression and localization of EIPR-1.**

(A) Representative images of animals expressing GFP under the promoter region upstream of the first gene (W09C5.1) of the operon where *eipr-1* is located. The image on the left shows expression in the hypodermis (skin) while the image on the right shows expression in the pharynx.

(B) Representative images of neurons expressing *eipr-1*::GFP under panneuronal (Left panel, *rab-3p*) and head cholinergic (Right panel, *unc-17Hp*) promoters. Scale bars: 5 μm.

(C) Representative images of neurons coexpressing *rab-3p*::*eipr-1*::GFP and GTP-bound RAB-2 (RAB-2QL). Scale bars: 5 μm.

**S5 Figure. EIPR-1 interacts with members of the GARP and EARP complexes.**

List of top hits from two independent experiments (A, B) performing mass spectrometry of a pulldown of rEIPR1::GFP in 832/13 cells. In A, we show the list of hits found after subtracting hits found in a GFP control pulldown. In B, we show the list of hits found after subtracting hits found in either the pulldown of CCCP1::GFP or RUNDC1::GFP. In both cases, EIPR1::GFP pulldowns were performed in parallel to controls. # seq = number of unique peptides from each protein. All proteins with two or more unique peptides are shown.

**S6 Figure. vps-51 mutants have defects in trafficking FLP-3::Venus.**

FLP-3::Venus fluorescence levels in the dorsal nerve cord of wild type and *vps-51* mutant strains. *vps-51* mutants have decreased fluorescence in the dorsal cord. Error bars = SEM; n = 8.

**S7 Figure. EARP, not GARP, is important for locomotion and trafficking dense-core vesicle cargos.**

(A). GARP mutants *vps-54*(*ok1463*) and *vps-54*(*ok1473*) do not have a reduced locomotion rate. Error bars = SEM; n = 10.

(B) *vps-50* acts in the same genetic pathway as *eipr-1* to control FLP-3::Venus trafficking. Left: representative images. Right: quantification. *vps-50* and *eipr-1* have reduced FLP-3::Venus fluorescence, but an *eipr-1*; *vps-50* double mutant does not have a stronger phenotype than either single mutant indicating that *eipr-1* and *vps-50* act in the same genetic pathway. A *vps-54* mutant does not have decreased axonal levels of FLP-3::Venus. ***, P<0.001 compared to wild type. Error bars = SEM; n =9-14.

(C) *vps-50* mutants do not have defects in trafficking synaptic vesicle cargos. SNB-1::mCherryfluorescence levels were measured in the dorsal nerve cord. ns, not significant, P>0.05 compared to wild type. Error bars = SEM; n =11-14.

**S8 Figure. VPS50 partially colocalizes with RAB2 and CCCP1.**

(A) Representative images of 832/13 cells expressing GFP::RAB2A and costained for endogenous VPS50. Scale bar: 10 μm.

(B) Representative images of 832/13 cells expressing CCCP1::GFP and costained for endogenous VPS50. Scale bar: 10 μm.

**S9 Figure. *eipr-1* and GARP/EARP mutants have reduced secretion of NLP-21::Venus.**

NLP-21::Venus fluorescence levels in the coelomocytes of the indicated strains. Like *rab-2* mutants, *eipr-1*, *vps-50*, and *vps-52* mutants show reductions in the accumulation of Venus fluorescence in coelomocytes that is approximately proportional to the decrease in axonal fluorescence seen in these mutants, suggesting that these mutants are not defective in dense-core vesicle release. Double mutants between *eipr-1* and *rab-2* or *vps-50* do not have stronger phenotypes than the single mutants, suggesting that these genes all act in the same dense-core vesicle cargo sorting pathway. **, P<0.01; ***, P<0.001 compared to WT. Error bars = SEM; n = 14-33.

**S10 Figure. *eipr-1* and *vps-50* mutants do not have enlarged lysosomes.**

Representative images of LMP-1::GFP fluorescence in coelomocytes of WT, *eipr-1*(*tm4790*) and *vps-50*(*ok2627*) mutant strains. Scale bar: 10 μm.

**S11 Figure. VPS-52 localizes normally in *eipr-1* mutants.**

Representative images of WT and *eipr-1*(*tm4790*) mutant neurons expressing VPS-52::tagRFP. Scale bars: 1 μm.

**S12 Figure. EIPR-1 interacts with SNAP29.**

EGFP-tagged rat EIPR1 or EGFP was expressed in 832/13 cells. Immunoprecipitation of EIPR1::EGFP pulled down more SNAP29 than untagged EGFP. IN: input; IP: immunoprecipitation.

**S1 Table. List of strains.**

**S2 Table. List of plasmids.**

